# Controlling Payload Heterogeneity in Lipid Nanoparticles for RNA-Based Therapeutics

**DOI:** 10.1101/2025.06.11.659145

**Authors:** Turash Haque Pial, Sixuan Li, Jinghan Lin, Tza-Huei Wang, Hai-Quan Mao, Tine Curk

## Abstract

Lipid nanoparticles (LNPs) are a leading platform for nucleic acid delivery, yet conventional assembly by mixing lipids with RNA yields particles with heterogeneous, bimodal payload distribution. Our transfection experiments demonstrate that heterogeneous siRNA distribution within LNPs significantly reduces gene knockdown efficiency. To systematically investigate the origin and extent of payload heterogeneity, we integrate coarse-grained molecular dynamics and kinetic Monte Carlo simulations with single-particle characterization via cylindrical illumination confocal spectroscopy and machine learning analysis. We find that the balance between RNA diffusion kinetics and lipid self-assembly dynamics is the dominant driver of payload heterogeneity. Leveraging this mechanistic insight, we show that (i) finely controlled turbulent mixing minimizes payload variance and increases the uniformity of RNA distribution without altering LNP size, and (ii) systematic adjustment of salt concentration and PEG-lipid content tunes RNA loading in a volume-dependent manner. Together, these results elucidate the self-assembly landscape of RNA-LNPs and provide actionable design principles for crafting more uniform, potent, and safer LNP-based nucleic acid therapies.

## Introduction

Nucleic acid-based therapeutics have revolutionized disease treatment, with lipid nanoparticles (LNPs) established as an effective delivery platform.^1–6^ The success of LNPs in FDA-approved vaccines, which deliver messenger RNA (mRNA), underscores their immense potential as a versatile delivery platform. Beyond vaccines, LNPs are being developed for various therapies, including small interfering RNA (siRNA)-mediated gene silencing,^7,8^ mRNA-based protein replacement,^9^ gene editing using mRNA and guide RNA,^10^ circular RNA for long-lasting protein expression in genetic disorder treatments,^11^ and DNA/RNA-based induction of immune responses against cancers and chronic infections.^12–15^ Despite this progress, a key gap lies in achieving precise control over how therapeutic payloads are distributed among individual LNPs as unwanted heterogeneous payload distribution can compromise therapeutic performance.^16–19^ Therefore, establishing a mechanistic understanding of payload heterogeneity and its effect on drug delivery is needed for optimizing nucleic acid-based delivery systems, and thereby de-risk and broaden the applicability of the LNP delivery platform.

RNA-LNPs are formed through a self-assembly process, where an aqueous solution of RNA is mixed with an alcoholic lipid solution containing ionizable lipids, PEGylated lipids, cholesterols, and helper lipids. Two key interactions primarily drive the self-assembly: the amphiphilic nature of the lipids, and electrostatic interactions between the negatively charged RNAs and the positively charged ionizable lipids. However, this assembly process often produces LNPs with heterogeneous compositions, including LNPs without RNA (“empty LNPs”) and LNPs containing multiple RNA copies, characterized by a bimodal RNA loading distribution^20^. This payload heterogeneity can significantly impact therapeutic outcomes, leading to toxicity or reactogenicity.^16,17^ A recent study has linked empty LNPs containing YSK13 lipids to liver toxicity, suggesting that the lipid component itself contributes to adverse effects.^18^ This highlights the importance of controlling the dose of administered RNA-LNPs to minimize toxicity, which necessitates precise RNA distribution within the LNPs. Researchers also found that the higher mRNA loading reduces transfection potency, likely due to their deviation from the optimal lipid/mRNA ratio and mRNA-mediated bleb-like structure formation.^21^ However, according to our knowledge, no study about potency is available for shorter or non-bleb forming RNAs like siRNA. Although less critical in vaccines, minimizing undesirable inflammation, improving potency, and enhancing drug delivery consistency are critical in long-term therapies such as gene delivery.

Formulating LNPs with desired physical properties, such as average size and RNA encapsulation efficiency, is straightforward, especially with techniques like microfluidics.^22–24^ However, controlling and measuring payload heterogeneity remains a significant challenge. Recently, we developed Cylindrical Illumination Confocal Spectroscopy (CICS), enabling analysis of mRNA, siRNA, or other payload characteristics at the single-particle level.^20,25^ This method uncovered substantial payload heterogeneity and a bimodal loading distribution, with up to 50% of LNPs being empty. While bimodal distribution is typically caused by thermodynamic phase separation, it may also stem from kinetic factors. The key question is whether thermodynamic or kinetic effects dominate. If the cause is kinetic, enhancing mixing could reduce heterogeneity. On the other hand, if the cause is thermodynamic, allowing the LNP configurations to equilibrate via enhanced mixing would likely increase heterogeneity. Thereby, identifying the precise cause of heterogeneity is crucial for optimizing strategies to obtain the desired RNA payload distribution.

In this study, we systematically explore the origin and extent of RNA payload heterogeneity and its effect on delivery efficiency. We investigate the formation of empty LNPs and demonstrate how to control their population. To this end, we first develop advanced multiscale computational techniques, employing molecular dynamics (MD) to model the initial stage of the RNA-LNP assembly process and kinetic Monte Carlo (kMC) to model the later stages, thus overcoming the challenge of limited simulation timescales. We investigate the effect of mixing rate, salt concentration (c_salt_), and PEGylation on the LNP size and payload distribution, and use a random-forest machine-learning method to analyze the effect of these features. Findings are validated using the CICS single nanoparticle detection experiments (Figure 1). Finally, we establish design principles to obtain a preferred LNP size and payload distribution.

**Figure 1.**
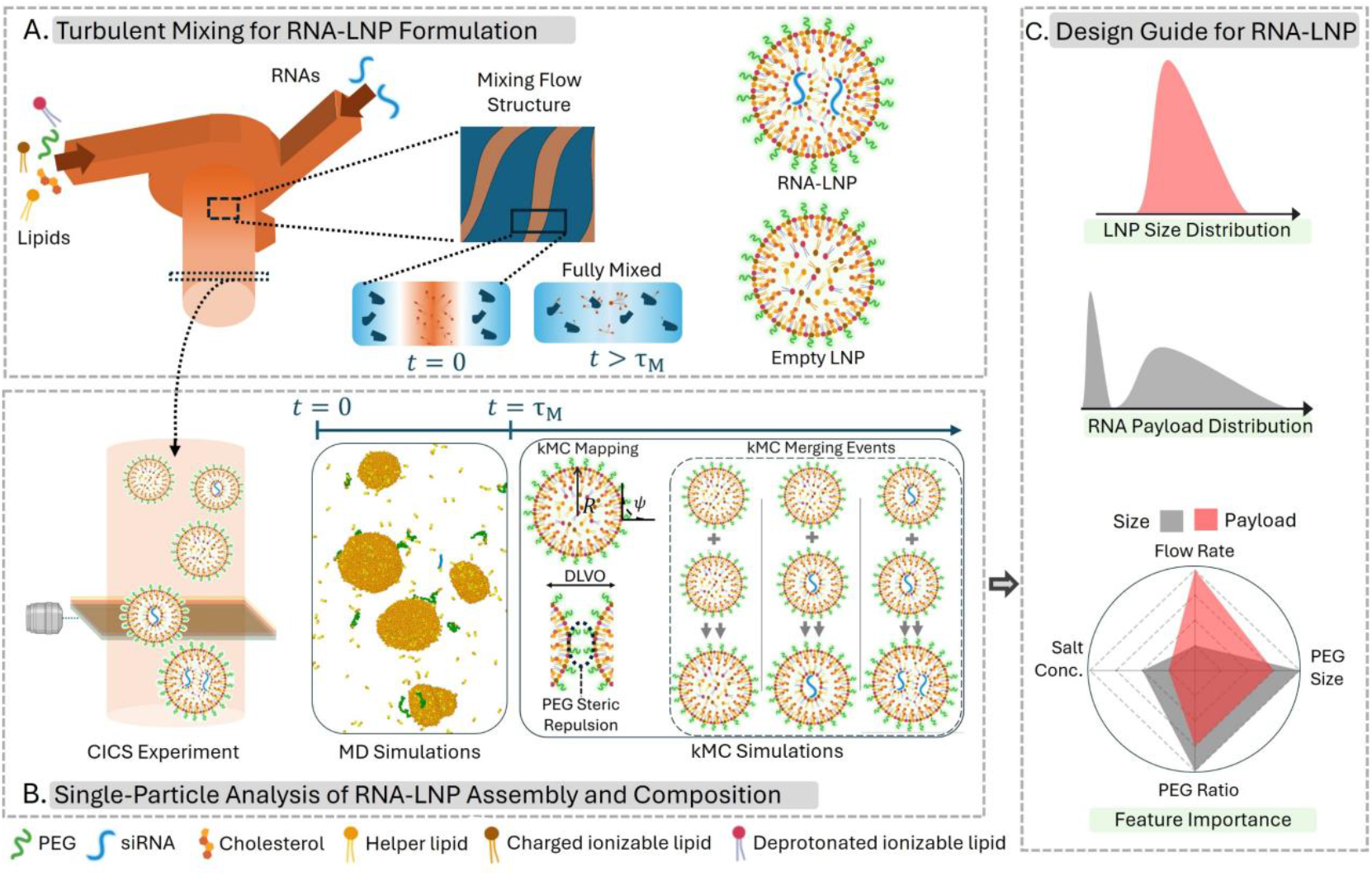
Overview of the RNA-LNP Self-Assembly Process: Experimental and Modeling Framework. (A) Schematic illustrating the self-assembly process of RNA-LNPs, where lipid-ethanol and RNA-aqueous solutions are mixed using a vortex mixer (left). Inlet flows generate turbulent inter-shearing layers, keeping lipids and RNAs separate until full mixing occurs due to diffusion on timescale τ_M_. This process resulted in RNA-loaded and RNA-free LNPs (empty-LNPs). (B) Cylindrical illumination confocal spectroscopy (left) is employed to analyze particle composition. Molecular dynamics (MD) simulations capture the initial stages of the assembly process (middle), whereas kinetic Monte Carlo (kMC) simulations are used for later stages (right). kMC incorporate charge regulation, Derjaguin-Landau-Verwey-Overbeek (DLVO) interactions, and PEG-PEG repulsion. Upon merging, LNPs update their composition and associated interactions. (C) kMC simulations provide detailed insights into RNA-LNP size distribution, payload distribution, and assembly dynamics. Coupling these simulations with machine learning enables the development of design rules for optimizing RNA-LNP formulations.

## Results

We utilized a benchmark lipid formulation based on the FDA-approved siRNA drug Patisiran.^26^ Typically, RNA-LNPs are formed by mixing an RNA-containing aqueous solution with a lipid-containing alcohol solution in turbulent setups such as T-junctions,^27^ confined impinging jets,^28^ or multi-inlet vortex mixers.^29^ We used a 3-inlet vortex mixer at flow rates of 10, 20, and 30 mL/min, with a siRNA-to-lipid flow rate ratio of 3:1. After mixing and subsequent dialysis (conducted 1-hour post-mixing), the siRNA-LNPs were analyzed using CICS (at 18 hours post-mixing), where each particle passes through a detector individually. Fluorescence signals from the detector enabled differentiation among LNP populations, categorizing them into siRNA-encapsulated LNPs, empty LNPs, and free unencapsulated siRNAs (Figure 1A-1B). This experimental system served as a reference for parameter selection and validation in our modeling approach.

### MD simulations reveal the origin of empty LNPs

To investigate the initial stages of the siRNA-LNP self-assembly process, we employed coarse-grained (CG) MD simulations. In turbulent mixing setups, the kinetic energy from the inlet flows breaks the flow jets into inter-shearing layers or turbulent eddies as the solutions enter the mixing chamber.^30^ To replicate these inter-shearing layers in simulations, we initially divided the system into distinct siRNA and lipid layers, maintaining a volume ratio 3:1 (Figure 2A(i)). Simulations reveal that during the mixing process, most lipids quickly aggregate to form small nanoparticles before significantly interacting with siRNA (Figure 2A, 2B). This early lipid aggregation leads to the formation of primarily empty LNPs. To our knowledge, this represents the first MD simulation that captures the RNA-LNP mixing process in a manner consistent with experimental mixing conditions.

**Figure 2.**
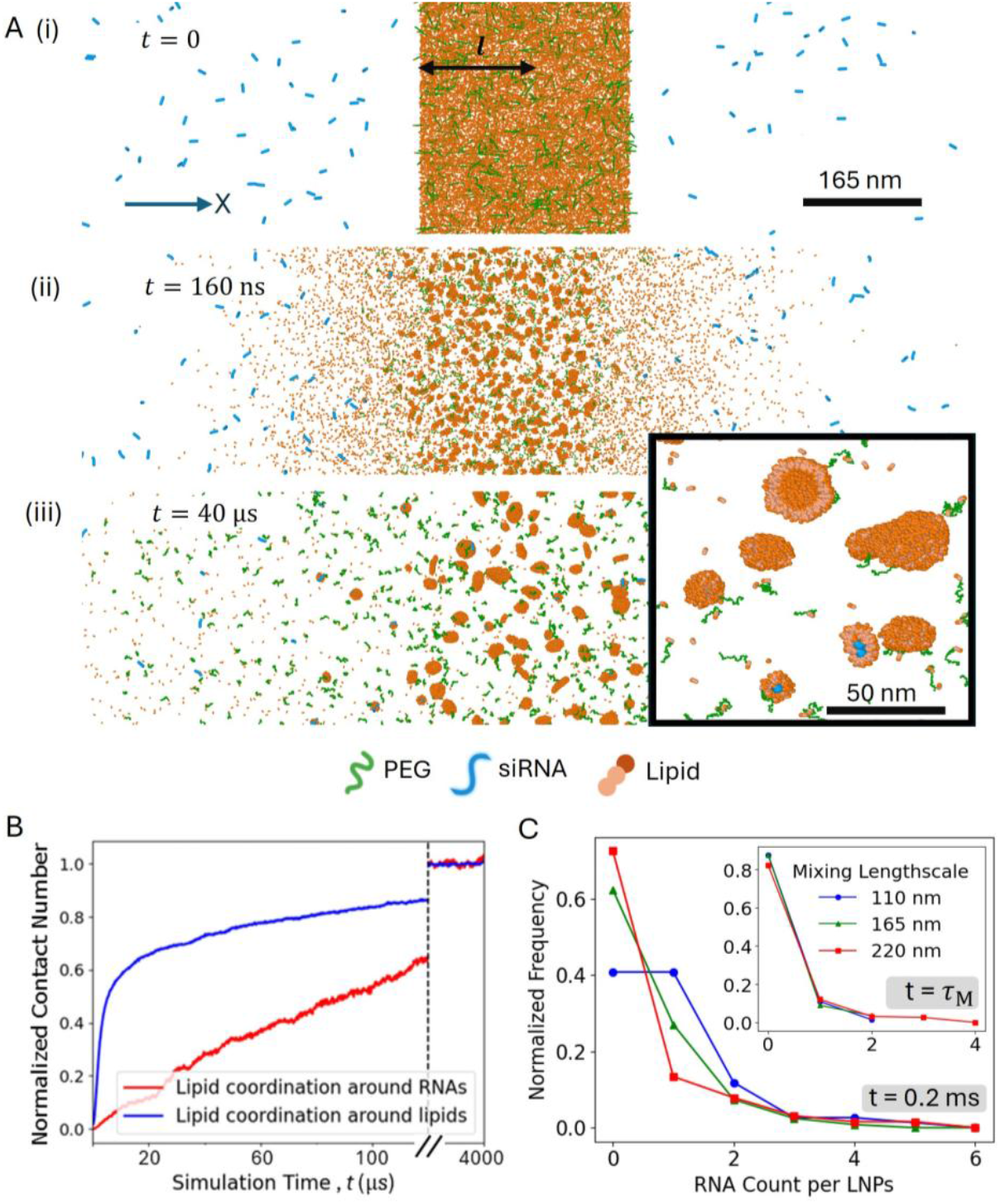
Molecular Dynamics (MD) Simulation of siRNA-LNP Assembly. (A) CG-MD simulation depicting the initial distribution of siRNA (blue), lipids (orange), and PEG (green) at (i) t=0, (ii) t=1600 τ, and (iii) t=40000 τ (τ= 2 ns). The characteristic mixing length scale, l (165 nm for these snapshots), is shown in black. Over time, lipids begin to aggregate and form small nanoparticles. Inset shows a zoom-in of the configuration, showing empty and siRNA containing LNPs. (B) Number of RNA–lipid, and lipid–lipid contacts normalized with the coordination number. (C) RNA payload distribution for different mixing length, l, at t=10^5^τ (or 0.2 ms). RNA distribution at mixing time scale τ_M_ =l^2^/2D, collapses to a master curve (inset). The τ_M_ values are about 25 μs, 57 μs, and 100 μs for l=110, 165, and 220 nm, respectively.

The thickness of inter-shearing layers can be controlled by adjusting the flow rate during siRNA-LNP formation.^30^ At higher flow rates, the increased kinetic energy leads to more intense breakdown of the flow and smaller turbulent eddies.^31^ In contrast, smaller flow rates may result in laminar flow, without any turbulent eddies. Following Ref. 30, we define the characteristic mixing length scale, *l*, as half the lipid layer thickness (Figure 2A). The distribution of siRNA on LNPs was analyzed at a simulation time of 100,000 *τ* (simulation time unit *τ* = 2 ns, yielding a total assembly time of 0.2 ms) at varying *l*. The results show that thicker inter-shearing layers (correspond to slower mixing rates in experiments) lead to a higher proportion of empty LNPs compared to thinner layers (Figure 2C). To quantitatively explain this behavior, we calculate the mixing timescale, *τ*_M_, for siRNA and lipids to diffuse across the mixing length scale, 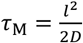 where *D* is the inter-diffusion constant (noting that in these simulations, *τ*_M_ values are about 25 μs, 57 μs, and 100 μs, respectively). Since individual siRNA molecules are much smaller than the LNPs, we approximate *D* ≈ *D*_siRNA_ (see supporting information SI, Figure S3). We find that at *τ*_M_ the loading distribution for different *l* collapse into a master curve (Figure 2C, inset). This demonstrates that the distribution of siRNA on LNPs is primarily governed by the diffusion timescale *τ*_M_, which is influenced by the thickness of the inter-shearing layers.

### kMC simulations and CICS experiments show the effect of mixing flow rates

Since MD simulations cannot capture the full growth dynamics of LNPs due to time scale limitations, we employ kMC simulations to model the aggregation of LNPs on timescales larger than the mixing time *τ*_M_. kMC simulations enable dynamic evolution of LNP growth^32^ and allow for the investigation of payload and size distributions. To initialize kMC simulations, we calculate the LNP radius *R*_0_ at mixing time *τ*_M_, using a mean-field coalescence theory of lipids (Figure 3A; see SI for derivation). We validated this approach by comparing the predicted sizes to those obtained from MD simulations. Based on MD simulation results (Figure 2C, inset), we set that at *τ*_M_ the fraction of empty LNPs *θ*_0_ = 0.83. In the turbulent mixing regime (flow rate *Q* ≥ 10 ml/min), the characteristic mixing time depends on the flow rate as *τ*_M_ = *b*_*Q*_/1.75 × *Q*^−1.5^, where *b*_*Q*_ = 1.3 × 10^3^ ms ^30,33,34^ which allows us to compare experimental observations with kMC simulations (the factor 1.75 accounts for diffusivity of siRNA relative to conditions in Ref. 30). A faster flow rate results in more turbulent mixing conditions with smaller *l* and shorter *τ*_M_ (Figure 2), and thus smaller size *R*_0_ (Figure 3A).

**Figure 3.**
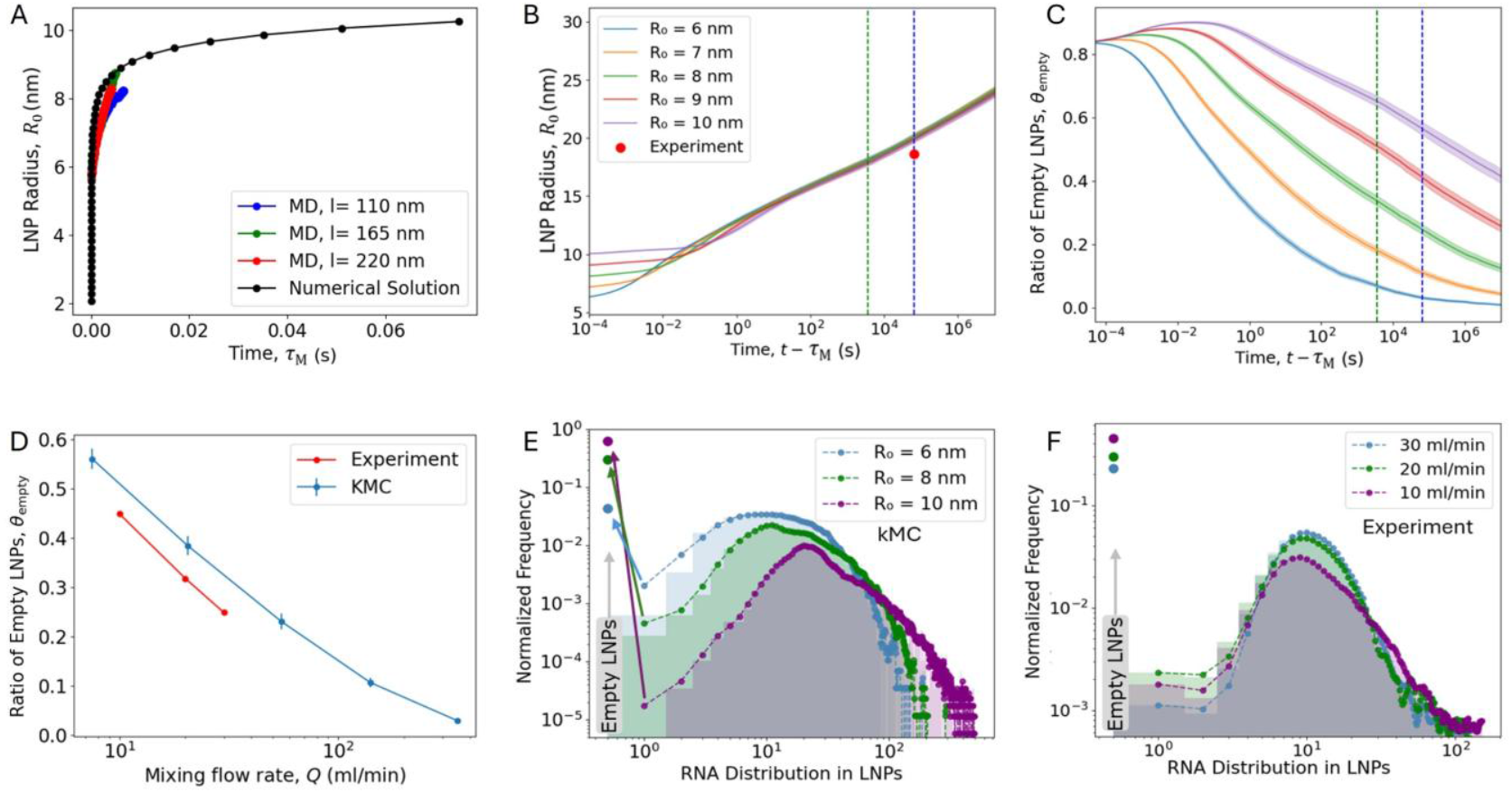
Effect of Mixing Flow Rate on LNP Size and Payload Distribution. (A) Theory prediction for LNP growth and MD simulation results providing the initial LNP radius input for kMC simulations. (B) Time evolution of the average LNP radius from kMC simulations. (C) Time evolution of the ratio of empty LNPs, defined as the fraction of LNPs without siRNA. The green vertical lines denote t = 1 h (post-mixing dialysis), and the blue lines denote t = 18 h (to match experimental measurement via CICS). (D) Ratio of empty LNPs as a function of mixing flow rate Q at t =18 h. (E-F) siRNA loading distribution at t =18 hours from kMC (E) and experiment (F). In both (E) and (F), normalized frequency distributions show a bimodal profile, with empty LNPs indicated on the left. The shaded regions show the raw histogram data, while the dashed lines show a moving average for visual clarity. Arrows in (E) help to show the bimodal nature of the loading distributions. Parameters: PEG MW is fixed at 2000 Da (RF=3.6 nm), PEG ratio 1.5% of total lipids, and the initial salt concentration csalt = 0.025 M.

Figure 3B illustrates the evolution of the average LNP radius *R* (note *R* is the bare LNP radius that does not include the PEG chains) in the kMC simulation. Surprisingly, particles reach an average radius of approximately 20 nm at 18 hours regardless of the initial size *R*_0_. This similarity in particle size for different *R*_0_ is also observed in experiment. Dynamic light scattering (DLS) intensity-weighted data reveals weak dependance of particle size on the mixing rate, with radii of 50.3 nm, 51.9 nm, and 52.5 nm at flow rates of *Q* = 10 ml/min, *Q* = 20 ml/min, and *Q* = 30 ml/min, respectively. This weak correlation is consistent with previous studies, which demonstrated that nanoparticle size decreases with increasing flow rates in laminar mixing regime but plateaus in the turbulent flow regime.^30,35^ Finally, according to our prior LNP experiments, DLS-derived average radii are about 2.8 times larger than those measured by cryo-TEM^20^ because DLS is skewed towards large particles, and we find that rescaling the DLS data agrees very well with the kMC simulations (Figure 3B).

To investigate the effects on LNP loading distribution, we analyzed the time-dependent changes in the ratio of empty LNPs *θ*_empty_ (Figure 3C), which is defined as the fraction of LNPs that contain no RNA, *θ*_empty_ = *N*_empty_/*N*_total_, with *N*_total_ the total number of LNPs and *N*_empty_ the number of LNPs containing no RNA. As LNPs merged into larger particles, the ratio of empty LNPs decreased overall. However, unexpectedly, the ratio increased in the early stages, which can be explained by the stronger electrostatic repulsion between empty LNPs, leading to LNPs containing siRNA to merge more quickly. Figure 3C demonstrates that reducing the initial LNP size significantly reduces the ratio of empty LNPs. Quantitative comparison between kMC results and experimental data at different mixing flow rates shows excellent agreement (Figure 3D).

While previous studies have shown the controllability of average nanoparticle size and drug loading efficiency, the distribution of internal components has remained largely unexplored. Our findings show that rapid mixing improves the uniformity/homogeneity of the internal composition within the nanoparticles. Surprisingly, this implies that the bimodal loading distribution results from mixing kinetics rather than thermodynamic phase separation. This is further supported by analyzing the entire siRNA loading distribution within LNPs (kMC data in Figure 3E, CICS data in Figure 3F). A larger initial LNP radius (smaller flow rate) results not only in more empty LNPs but also increases the number of LNPs with excessive siRNA (longer distribution tail), indicating highly heterogeneous encapsulation and a broader bimodal distribution. The agreement between kMC and experimental data (Figure 3B, D–F) is remarkable given that the kMC model contains no fitting parameters. This strongly suggests that the kMC model captures the relevant physical processes and can be used as an independent tool to predict LNP assembly.

### A more uniform payload distribution improves delivery efficiency in vitro

To investigate the impact of payload distribution uniformity on delivery efficiency, we performed in vitro transfection assays using LNPs generated at flow rates of 1 mL/min (not covered in the kMC calculations), 10 mL/min, and 30 mL/min. Increasing the flow rate reduced both the average siRNA payload and the ratio of empty LNPs, indicating that higher flow rates yield LNP populations with more uniform and efficient siRNA encapsulation (as shown in Figure 3 E, F; high flow rate leads to narrower payload distribution, see also Figure 6D). Note that LNPs formed at the lowest flow rate (1 mL/min, corresponding to the laminar mixing regime) were larger in average size (62.0 nm in radius, compared to 51.0 nm and 52.5 nm for 10 ml/min and 30 ml/min, respectively), which resulted in LNPs with a higher average siRNA payload per particle.

At a high siRNA dose (100 ng), all formulations achieved near-complete knockdown of GFP expression in GFP+ cells (Figure 4A). This indicates that at saturating concentrations, the influence of payload distribution becomes negligible. However, at lower doses, the differences in knockdown efficiency diverged: LNPs prepared at 30 ml/min consistently outperformed those produced at lower flow rates. These particles, with a more uniform payload distribution and fewer empty LNPs, achieved greater GFP silencing and lower mean fluorescence intensity (MFI, Figure 4B) than the more heterogeneous populations from 1 and 10 ml/min.

**Figure 4.**
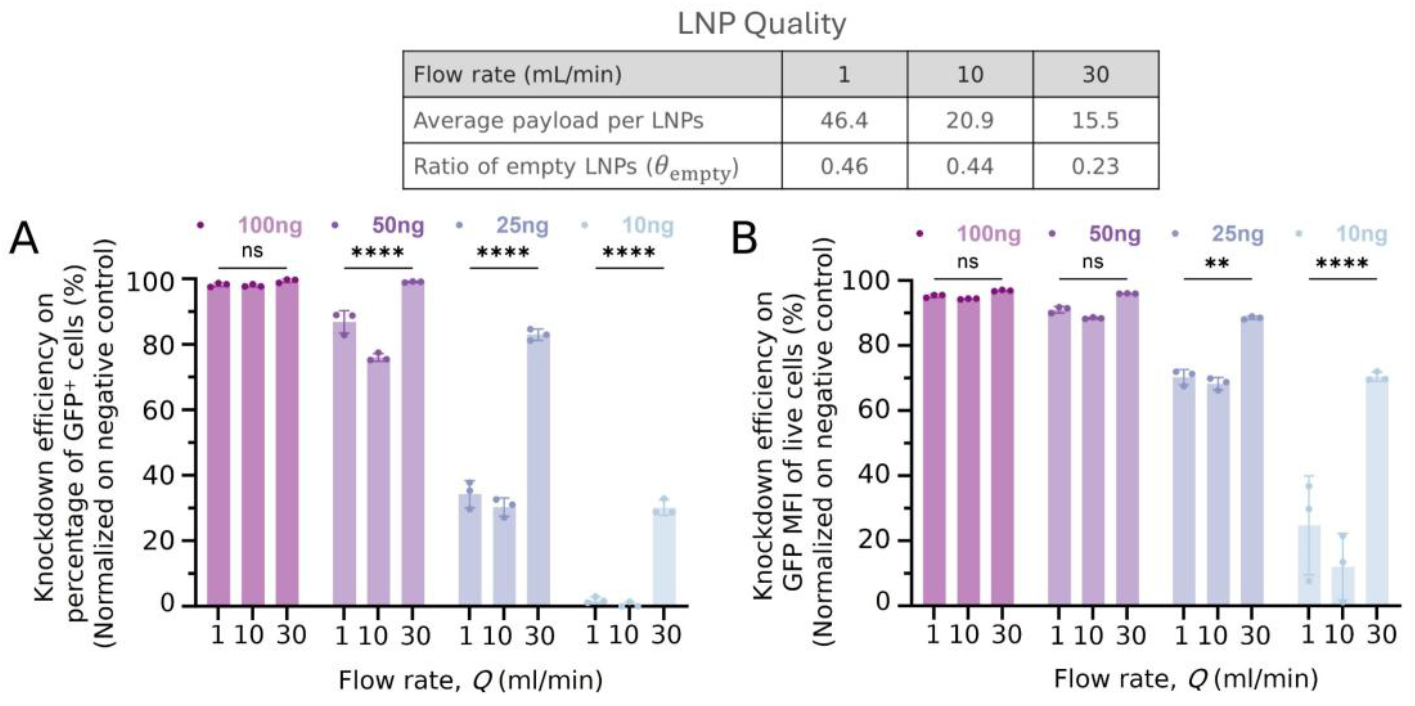
Effect of payload heterogeneity in transfection efficiency. The top table summarizes siRNA-LNP quality metrics, illustrating improved payload uniformity and reduced fraction of empty LNPs at higher flow rates. (A, B) Knockdown efficiency of GFP expression in vitro (normalized to negative control) as a function of flow rate used during LNP formulation and siRNA dose. Higher flow rates produce more uniform LNPs with a reduced ratio of empty LNPs (top table), enhancing knockdown efficiency at low siRNA doses. At high doses, all formulations achieve near-complete GFP silencing.

These results demonstrate that improved knockdown efficiency at lower doses correlates strongly with the uniformity of siRNA payload distribution within the LNP population. At lower flow rates, the broader payload distribution negatively affected knockdown efficiency, particularly at subsaturating doses. Importantly, the encapsulation efficiency was greater than 90% across all conditions, ensuring a consistent total number of delivered siRNA across equivalent doses. Therefore, the observed differences in knockdown efficiency can be attributed exclusively to variations in payload distribution among the LNP populations rather than total siRNA content.

### kMC simulations show the effect of PEG size and salt concentration

Along with mixing setup, PEG molecular weight (MW) is crucial in LNP assembly, stability, and delivery efficiency.^36–38^ PEG forms a hydration shell that prevents LNP aggregation and extends circulation time. LNPs with shorter PEG (e.g., 750 Da) show little difference from PEG-free LNPs. However, longer PEG chains (over 5000 Da) can hinder cellular uptake and endosomal escape, key steps in delivering therapeutic agents.^38^ Therefore, understanding the intermediate range (1000-5000 Da) is critical for optimizing stability and RNA delivery.

We investigated the effect of PEG MW by adjusting the Flory radius, *R*_F_ = a*n*^3/5^ in PEG mediated repulsion calculation (a=0.37 nm is the monomer size and *n* the degree of polymerization).^39,40^ Figure 5A shows that increasing the PEG MW slows LNP growth, creating an inverse relationship between PEG MW and LNP size. This well-known effect^25,37^ is explained by the increased steric barrier *E*_PEG_ resulting from a larger *R*_F_ (see Eq. 2 in the Methods section). On the other hand, increasing PEG monotonically increases the ratio of empty LNPs (Figure 5B), reduces the normalized frequency of RNA loading (Figure 3C) and yields a broader distribution. A similar effect is expected when increasing the PEG fraction, as both variables influence the PEG mediated repulsions(see Eq. 2 and Ref. 20).

**Figure 5.**
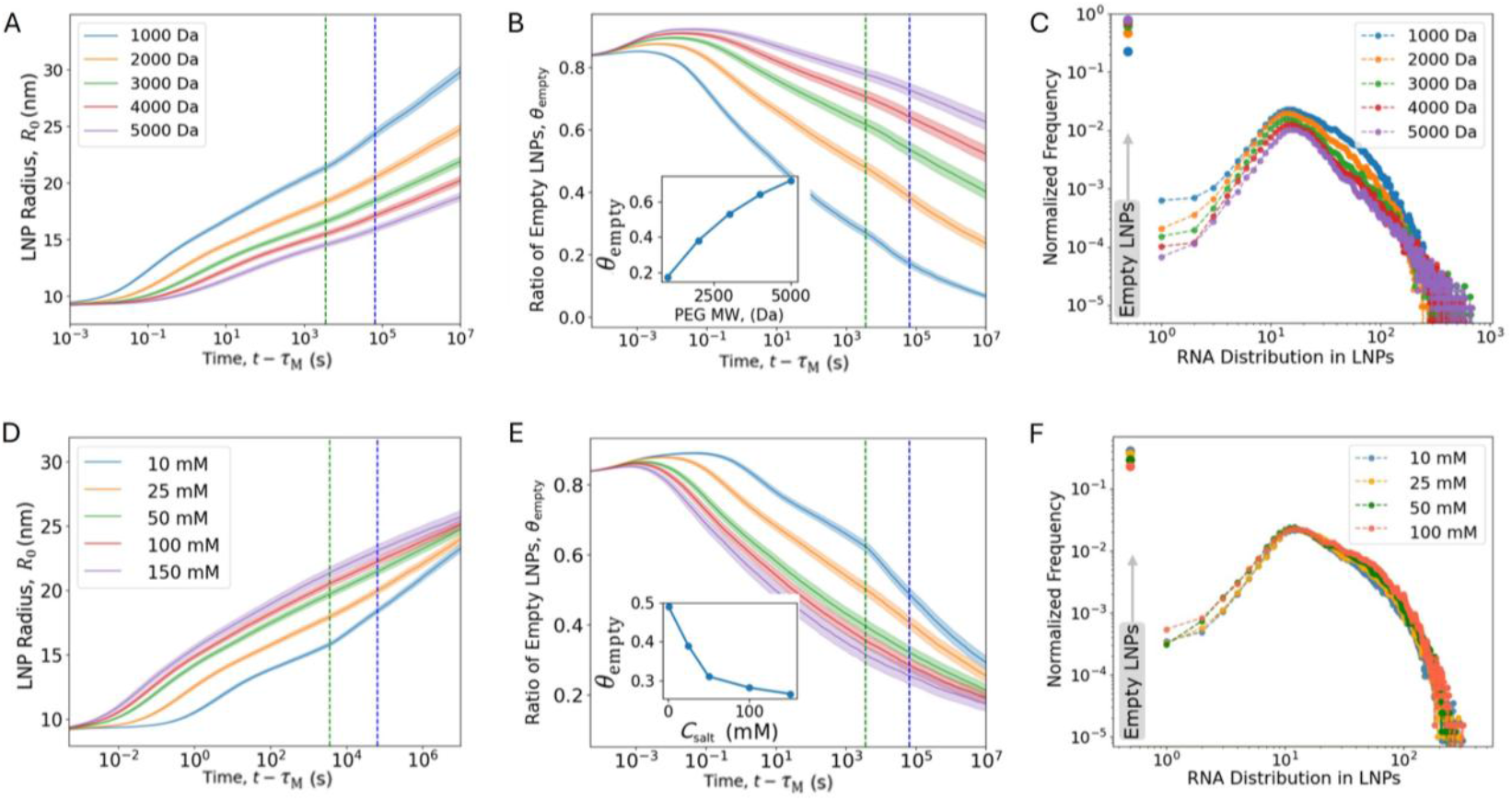
Effect of PEG Size and Salt Concentration on LNP Size and Payload Distribution. (A) LNP growth and (B) ratio of empty LNPs from for varying PEG MW. The green vertical line denotes t = 1 h (post-mixing dialysis), and the blue line denotes t = 18 h (to match experimental measurement via CICS). Inset in (B) shows the ratio of empty LNPs at 18 hours. (C) siRNA loading distribution at t = 18 h. (D) LNP growth and (E) ratio of empty LNPs for different c_salt_. Inset in (E) shows the ratio of empty LNPs for varying salt concentration at 18 h. (F) siRNA loading distribution at t = 18 h. Parameters: Initial LNP size is fixed at 9 nm, PEG ratio is 1.5% of total lipids, in (A, B, C) initial salt concentration c_salt_ = 0.025 M, while in (D, E, F) PEG MW = 2000 Da (R_F_ = 3.6 nm).

**Figure 6.**
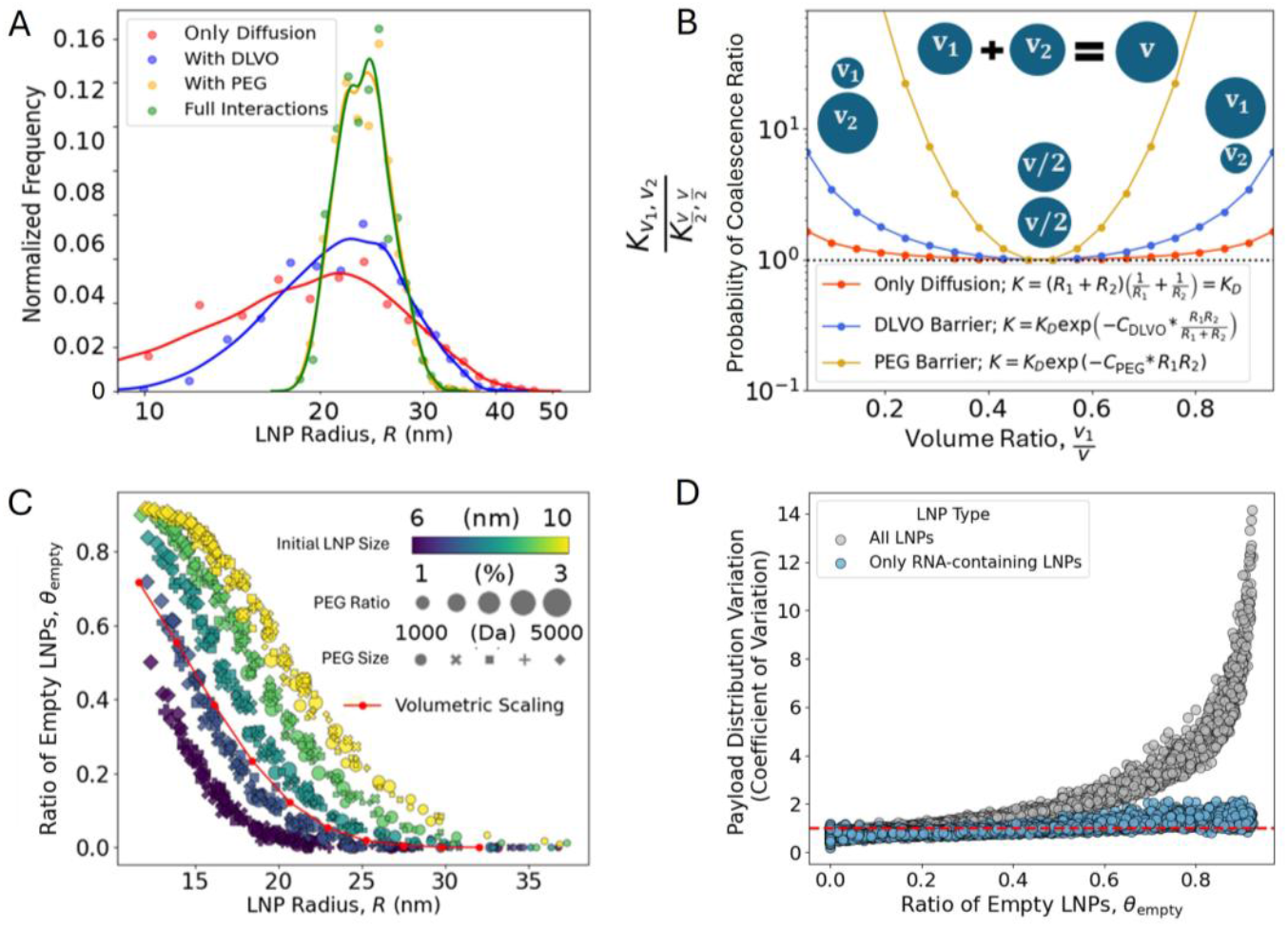
Analysis of Size Polydispersity and Payload Distribution Heterogeneity. (A) LNP size distributions from kMC simulations using different aggregation kernels. Kernel density estimation (KDE) lines are overlaid to highlight the distribution profiles. (B) Aggregation bias expressed as the ratio 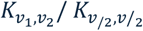, i.e., the probability of coalescence between an LNP of volume v_1_ and one of volume v_2_, relative to the probability of coalescing two LNPs of volume v/2, to form a larger LNP of volume v. This illustrates size-selective merging behavior. (C) Correlation between the ratio of empty LNPs and the average LNP radius at t = 18 h from kMC simulations. The red line indicates a volumetric scaling, with R_0_ = 10 nm and θ_0_ = 0.83. (D) Correlation between the ratio of empty LNPs and the pooled coefficient of variation (CV) of siRNA payload distribution across LNP populations. The red dashed line shows the reference value CV = 1.

In addition to PEG, salt concentration c_salt_ plays a key role in the self-assembly of LNPs by modulating electrostatic interactions. We explore an experimentally relevant range of *c*_salt_, from 10 mM to 150 mM. kMC simulations reveal that increased *c*_salt_ accelerates aggregation by reducing electrostatic repulsion between cationic LNPs, as shown in Figure 5D. Beyond 50 mM, *c*_salt_ yielded similar aggregation kinetics, due to neutralization of surface charge and reduction of electrostatic surface potential. Additionally, the faster aggregation at higher *c*_salt_ reduces the proportion of empty LNPs (Figure 5E and inset). Changes in *c*_salt_ yield a similar effect on siRNA distribution (Figure 5F) as changes in the PEG MW (Figure 5C), although the effect of salt is less pronounced compared to PEG.

### Analysis of LNP polydispersity and payload distribution

The above findings, combining kMC simulations and experimental data, highlight the complex interplay between mixing behavior, steric repulsion, and electrostatic interactions in LNP coalescence and growth. To further unravel precisely what factors determine the LNP heterogeneity, we isolate and analyze how different parameters affect LNP size and payload distribution by selectively deactivating the contributions from PEG (Eq. 2) and electrostatic and van der Waals (DLVO) interactions (Eq. 3) in kMC. Excluding both PEG and DLVO interactions results in LNP coalescence driven solely by diffusion, leading to a very broad LNP size distribution (Figure 6A). Minimizing size polydispersity is crucial for improving the performance, stability, and safety of LNPs.^1^ Reintroducing either DLVO or PEG interactions narrows the distribution, with PEG having a significantly stronger effect.

To understand why PEG exerts greater influence, we examined the aggregation kernel *K* and the associated fusion energy barriers which reveal size-selective merging behavior. The PEG contribution to the fusion barrier between two spheres with radii *R*_1_ and *R*_2_ is *E*_PEG_ = *C*_PEG_*R*_1_*R*_2_, and for DLVO interactions it is 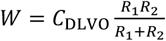 (Eqs. 2 and 3). Here, *C*_PEG_ and *C*_DLVO_ are simplified multiplicative factors, derived using typical values in Eqs. 2 and 3 (see Figures S7 and S8). In Figure 6B, we plot the aggregation bias ratio 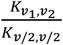, representing the probability of coalescence between an LNP of volume *v*_1_ and with another LNP of volume *v*_2_, relative to the probability of merging two LNPs of volume *v*/2 to form a larger LNP of volume *v*. This ratio reveals that the probability of merging a smaller LNP with a larger one is higher than the probability of merging two LNPs of similar sizes. This behavior promotes a narrower size distribution by: (1) reducing the number of smaller LNPs and (2) limiting the formation of very large LNPs. We find that this effect is stronger for PEG compared to DLVO, as the DLVO fusion barrier increases exponentially with LNP radius, whereas the PEG fusion barrier increases as a squared exponential. Thus, PEG steric repulsion is the main factor that determines the LNP size and size polydispersity, with DLVO electrostatic repulsion having a smaller effect.

Having analyzed the size polydispersity, we now turn to payload distribution. Investigating the correlation between LNP size and the payload distribution, and whether these properties can be independently controlled, is crucial for optimizing LNP formulations. Surprisingly, we find that the PEG properties have no influence on the ratio of empty LNPs *θ*_empty_ (see overlapping symbols in Figure 5C), instead *θ*_empty_ is determined solely by the initial and final LNP size. The ratio of empty LNPs *θ*_empty_ and the average LNP radius *R* behave in a manner close to volumetric scaling: 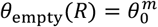, where *θ*_empty_(*R*) is the fraction of empty LNPs at radius *R, θ*_0_ the fraction of empty LNPs at *R*_0_, and *m* = (*R*/*R*_0_)^3^ the average number of merging events to grow an average particle size from *R*_0_ to *R*. The slight deviation of kMC results compared to volumetric scaling (Figure 6C) is attributed to the non-constant aggregation kernel, as the probability of merging is not equal for all LNPs. Thereby, mechanistically, changes in PEG or salt concentration directly affect the LNP size and size polydispersity, but do not directly affect the payload distribution (see overlapping symbols in Figure 6C), instead their effect on the payload distribution is indirect, a secondary effect due to the change in the average size *R*.

In addition to understanding the fraction of empty particles, it is also important to predict the shape of the full loading distribution (Figures 3E,F and 5C,F). We calculate the pooled coefficient of variation (CV) of siRNA loading distributions across LNP populations (Figure 6D). The CV, defined as the standard deviation divided by the mean, quantifies the payload variability in a population. Two metrics were computed: (1) the total CV across all LNPs, including empty particles, and (2) the CV restricted to non-empty LNPs only. We observed that the heterogeneity in payload distribution increases monotonically with the fraction of empty LNPs, even for non-empty LNPs. This suggests that inefficient mixing not only result in more empty particles but also lead to greater variability in payload among loaded LNPs. For instance, this effect is clearly illustrated in Figure 3E,F where the peak frequency decreases and the distribution tail broadens, indicating higher payload variation. All data points for all different parameters fall on two master curves in Figure 6D, implying that the shape of the full loading distribution is highly correlated to the fraction of empty particles. Together, the results in Figure 6 establish a foundational understanding of the underlying correlations, offering a framework for decoupling and independently tuning LNP size and RNA loading distributions.

### Machine learning elucidated design rules

To identify and quantify the most influential factors affecting LNP size and payload distribution, we performed a feature importance analysis using a random forest-based ensemble machine learning approach^41^. Feature importance quantifies how much each input variable (feature) contributes to output predictions. Our input parameters space includes: *R*_0_ (6 to 10 nm), PEG MW (1000 to 5000 Da), PEG ratio of lipids (1 to 3%), and c_salt_ (10 to 150 mM). The analysis reveals that the initial LNP size or mixing flow rate has a negligible effect on the final average LNP size (Figure 7A), whereas the PEG molecular weight (MW) and the PEG ratio play the most significant roles in determining LNP size. c_salt_ can also influence LNP size, but its effect is constrained by the upper limit (charge neutralization for c_salt_ > 50 mM), as well as a relatively weaker effect compared to PEG (as seen in Figure 6B). Interestingly, the flow rate emerges as the most critical factor controlling the ratio of empty LNPs and the payload distribution, despite having minimal influence on the final LNP size (Figure 7A).

**Figure 7.**
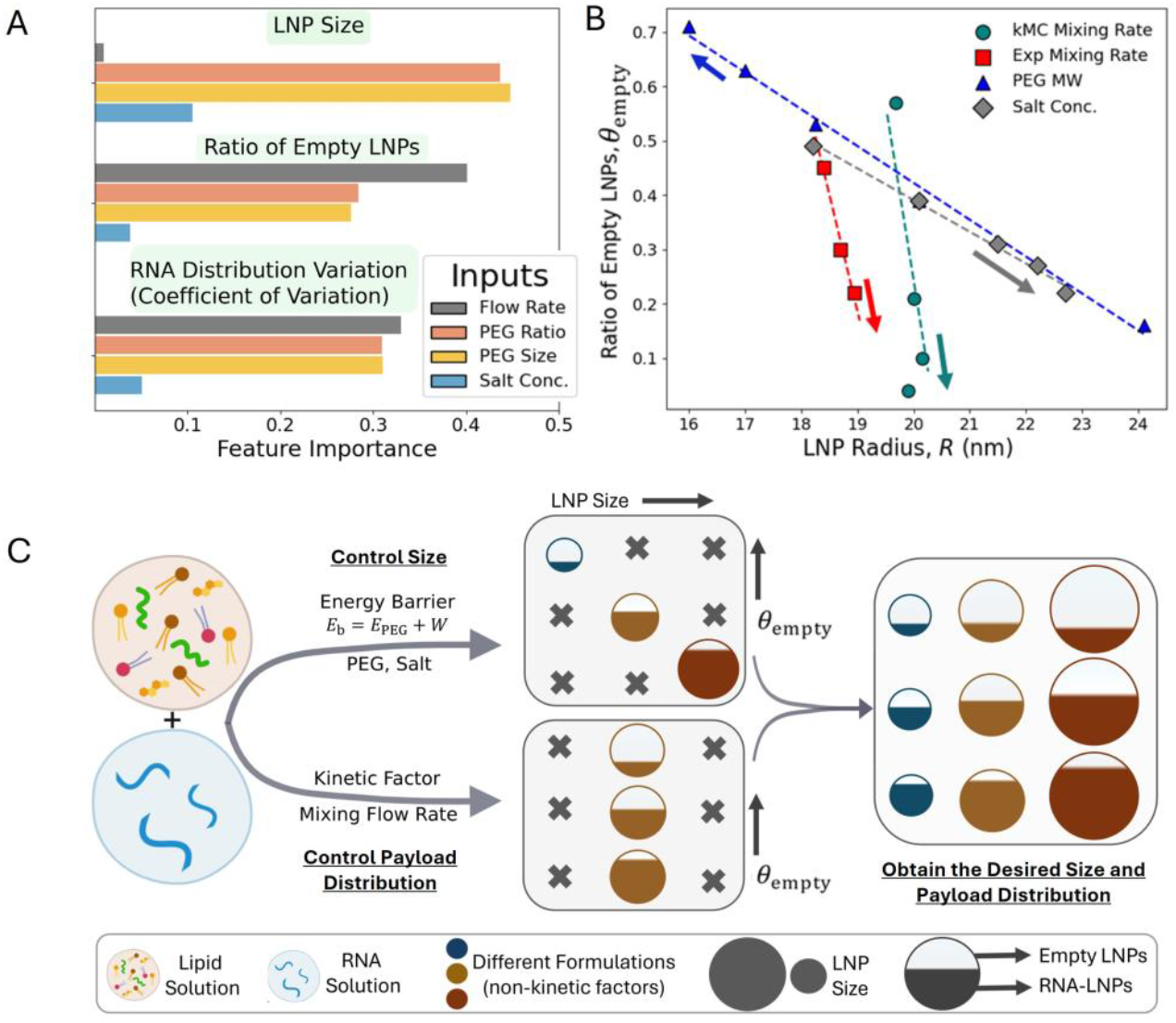
Design Guide for LNP Size and Payload Distribution. (A) Random-forest feature importance analysis highlights the relative importance of different parameters in controlling LNP size and payload. The corresponding SHapley Additive exPlanations (SHAP), which shows positive and negative correlation of features with model predictions is shown in SI Figure S9. (B) Mixing rate strongly affects the ratio of empty LNPs θ_empty_, but only weakly affects the LNP size R, in both kMC and experiment. Volumetric trend is observable for PEG and salt concentration (Figure 5). (C) Schematic demonstrating how the non-kinetic (energy barrier) and kinetic factors influence the LNP size and payload distribution. LNP energy barrier affects both the LNP size and payload distribution, whereas kinetic measures (e.g. mixing flow rate) independently control only the payload distribution. Crosses indicate inaccessible regions that cannot be achieved using either kinetic or non-kinetic tuning alone. By integrating both kinetic and non-kinetic factors, a wide range of LNP sizes and payload distributions can be achieved.

The LNP size and loading distribution data from Figures 3 and 5 are summarized in Figure 7B. Changes in mixing rate yield a nearly vertical slope on the *θ*_empty_–*R* parametric plot, observed in both computational and experimental data, highlighting the ability to independently control payload distribution and LNP size. In contrast, the gradual slopes for PEG and salt indicate a strong correlation between LNP size and payload distribution, showing a volumetric trend.

## Discussion

Kinetic Monte Carlo simulations and experimental results together demonstrate that the size and payload distribution of LNPs can be controlled by utilizing energetic and kinetic factors (Figure 7A, B). These insights can be used to produce LNPs with controlled size and payload distribution (Figure 7C). The molecular weight of PEG, the PEG ratio, and salt concentration all contribute to the LNP fusion barrier (Figure 6), thereby determining the final LNP size. Adjusting these non-kinetic parameters allows for precise control of the LNP size, while their effect on the fraction of empty particles *θ*_empty_ and the loading distribution is indirect; it is a secondary effect due to the change in LNP size. In contrast, the mixing flow rate serves as a kinetic control parameter that dictates the relative assembly of lipid-lipid and RNA-lipid, enabling regulation of the RNA payload distribution independently of particle size. By integrating both types of control parameters, it becomes possible to produce LNPs with desired combination of size and payload distribution. Such precision is critical because LNP size strongly affects therapeutic delivery and cellular uptake, while homogeneity in RNA loading is important for maximizing potency and ensuring safety.

Our integrative computational framework combining molecular dynamics with kinetic Monte Carlo and a surrogate model for electrostatic interactions offers new mechanistic insights and practical design rules. This computational framework is generalizable to a broad class of self-assembly systems in solution. Within the kMC model, the chemical characteristics of lipid components can be systematically tuned by adjusting parameters such as pKa and molecular mass. Moreover, key experimental variables, including the duration of dialysis, total assembly time, and the ratio or amount of different molecular components, are explicitly controllable in the simulation. The framework is also adaptable to diverse payloads, provided their mass and charge properties are properly defined. The simulation results show very good agreement with experimental observations, supporting the validity of the approach.

Despite the broad applicability of our LNP design rules, opportunities remain to extend the model and will shape future work. First, all simulations and experiments use a 22-bp siRNA payload. Larger cargos, such as >3 kb mRNA or plasmid DNA diffuse far more slowly. Extending the model to capture the interdiffusion and encapsulation of larger nucleic acids is an important next step, although we anticipate that the core design principles will remain the same. Second, the current parameter space; PEG 1–3 mol%, PEG MW 1–5 kDa, and 10–150 mM salt concentration covers common but not exhaustive formulation conditions. Systematically exploring higher PEG fractions, alternative buffer systems, broader flow-rate ratios, and a range of stoichiometry ratios will enhance the utility and generality of the proposed design map. This enhancement will extend the framework to other clinically realistic LNP formulations, accelerating the rational design of safer, more potent nucleic-acid therapeutics.

## Conclusion

In summary, through the integration of CG-MD simulations, kMC simulations, single-particle spectroscopy, machine learning, and transfections experiments, we have developed a comprehensive understanding of the siRNA payload distribution and its effect on delivery efficiency. We found that increasing homogeneity of siRNA distribution in LNPs is significantly improves drug delivery efficiency, especially at subsaturating doses. Our simulation model spans a wide range of time scales from microseconds to days, sufficient to capture experimentally relevant dynamics and heterogeneities. We found that the origin of empty LNPs and bimodal payload distribution is kinetic rather than thermodynamic. Bimodal RNA loading is caused by fast self-assembly of empty LNPs compared to the relatively slow diffusive mixing timescale. Rapid turbulent mixing of lipids and RNA significantly reduces empty LNPs and narrows the siRNA payload distribution without altering LNP size. While PEGylation and salt concentration also affect payload distribution, they do so only indirectly by changing the average LNP size. Additionally, kMC data shows that controlling the fusion barrier through PEG modification, rather than electrostatic stabilization alone, is crucial for achieving a narrow size distribution. These findings provide fundamental understanding of the LNP assembly process and offer essential design principles for optimizing LNP-based RNA therapeutics, aiming to enhance both their efficacy and safety. Additionally, the computational framework that we developed is freely accessible and can be extended to other therapeutic components, such as mRNA, and other nanoparticles, offering a powerful tool for efficiently screening nanoparticle formulation parameters.

## Methods

### Single particle analysis of siRNA LNP on CICS

For the single nanoparticle analysis of siRNA LNP, the LNPs use the same formulation stoichiometry as used in simulations, with DLin-MC3-DMA (MedKoo Biosciences, Cat.#555308), DSPC (Avanti Polar Lipids, Cat.#850365), cholesterol (Sigma-Aldrich, Cat. #C8667), DMG-PEG2000 (NOF America, Cat# GM020) at a molar ratio of 50:10:38.5:1.5 dissolved in 100% ethanol and RNA in 25 mM sodium acetate buffer at pH 4.0. For the fluorescence detection, the LNP sample was prepared with fluorescently labeled DSPC-TopFluor (Avanti Polar Lipids, Cat. #810281) substituting 10% DSPC lipid, Cy5 labeled DSPE PEG2000-N-Cy5 (Avanti Polar Lipids, Cat. #810891) substituting 33.3% PEG lipid and 100% siRNA-Cy3 (Sigma-Aldrich, Cat. #SIC003). The LNP samples were formulated by a 3-inlet vortex mixer^42^ at a total flow rate of 1, 10, 20, and 30 mL/min with a lipid to RNA flow rate ratio of 1:3. The final siRNA concentration after LNP formulation was 20 μg/mL. The sample was then (at 1 h) dialyzed against 1x phosphate buffered saline (PBS) buffer at pH 7.4 for 12 h under 4 °C using dialysis tubes with a molecular weight cutoff (MWCO) of 3,500 Pur-A-Lyzer dialysis kit, Sigma-Aldrich, Cat. #PURD35050). Following dialysis, the LNPs were analyzed by <u>C</u>ylindrical Illumination <u>C</u>onfocal <u>S</u>pectroscopy (CICS).^43^ The instrumentation and single-nanoparticle analysis have been developed in our previous work and are described in detail.^20,25^ Briefly, LNP samples were diluted sufficiently to allow for single particles or siRNA molecules to go through the detector one at a time. The 3-color fluorescence signals of each event were used to differentiate LNP populations into siRNA-encapsulated LNPs, empty LNPs, and free unencapsulated siRNAs. By applying the fluorescence deconvolution to the fluorescence distributions of siRNA encapsulated LNPs against a baseline of single siRNA run separately, the siRNA payload distribution on the single LNP level was obtained (see Figure S1).

### In vitro transfection

LNPs were prepared as described above, and Silencer™ GFP (eGFP) siRNA (Thermo Fisher Scientific, MA, USA) was encapsulated into the particles. The resulting LNPs were dialyzed against 1× PBS (pH 7.4) overnight at 4 °C. The final siRNA concentration was 20 µg/mL. B16F10-GFP cells were seeded in 96-well plates at 10,000 cells per well. On the following day, cells were treated with LNPs at doses of 100 ng, 50 ng, 25 ng, or 10 ng siRNA per well. After 48 h, cells were harvested and analyzed by flow cytometry using an Attune NxT instrument (Thermo Fisher Scientific, MA, USA).

### Molecular dynamics simulations

We modified Cooke and Deserno 3-bead lipid model^44^ by introducing attractive head-to-head interactions to promote nanoparticle formation (see Figure S2). Strength and cut-off values for these attractive head-to-head interactions are provided in the SI. siRNA was included at a nitrogen-to-phosphate (N/P) ratio of 6, and 1.5% of the lipid was PEGylated, using the same stoichiometry as the experimental setup. The 22 base-pair siRNA and 2000 Da PEG were represented with a bead-spring model. To match experimental conditions, 50% of the lipids carried a positive charge, interacting with the negatively charged siRNA via Debye–Hückel interactions at a salt concentration of 0.025 M.^45^ Since, solvent mixing (e.g., ethanol dilution into the aqueous solution) occurs much faster (as ethanol diffusion coefficient is 2 orders of magnitude greater than siRNA) relative to lipid and RNA mixing, we consider that ethanol and water are fully mixed. Simulations were conducted using a Langevin thermostat ensuring accurate diffusion behavior of RNA and lipids (see SI Figure S3). All simulations were performed using LAMMPS,^46^ and OVITO was used for visualization.^47^ RNA–lipid and lipid–lipid contacts (Figure 2B) were counted within the first valley of the radial distribution functions (see SI, Figure S4).

### Kinetic Monte Carlo simulations

We consider that during mixing time *τ*_M_, most lipids form spherical nanoparticles and that after *τ*_M_, the solution is well-mixed. Consequently, the rate per unit concentration at which two spherical nanoparticles with radii *R* _*i*_and *R*_*j*_ collide due to diffusion is 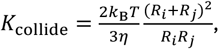^48^ with *k*_B_ the Boltzmann constant, η the viscosity, and *T* the absolute temperature. Assuming Arrhenius behavior, the rate at which two specific LNPs in a volume *V*_system_ merge is

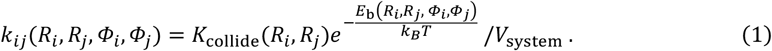

Here *E*_*b*_ is the fusion barrier between two LNPs with compositions *Φ*_*i*_, *Φ*_*j*_ (volume fractions of different components in LNPs); *E*_b_ is modeled by the summation of PEG repulsion (*E*_PEG_) and Derjaguin-Landau-Verwey-Overbeek (DLVO) interaction (*W*),^49^ *E*_b_ = *E*_PEG_ + *W*. We consider that PEG chains are mobile on the LNP and for the two particles to merge the PEG must be excluded from the interaction area, thus, the free-energy barrier can be calculated as an entropic penalty to exclude PEG chains from the interaction area,

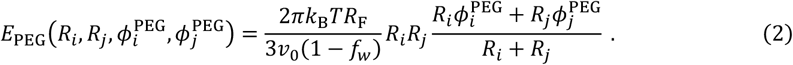

Here *f*_*w*_=0.2 is the water volume fraction in LNPs^20,50^, *v*_0_ the volume per lipid, *R*_F_ the Flory radius of PEG determined by the PEG size,^39,40^ and 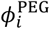 the PEG fraction in particle *i*. The DLVO interaction energy is a sum of van der Waals and electrostatic interactions^49^:

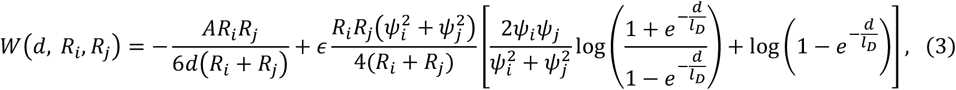

with *A* is the Hamaker constant, *l*_D_ the Debye screening length, *ψ* the surface electrostatic potential, *ϵ* the permittivity of solvent, and *d* the surface-to-surface distance between LNPs. The average time for a fusion event to occur is 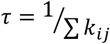, where the sum proceeds over all LNP pairs in the system. In the kMC implementation, a fusion event is chosen randomly according to the rate *k*_*ij*_, with the time increment determined by *τ*. Each simulation was initialized with 2500 LNPs, expect for those in Figure 5A where simulations start with N = 2000 LNPs and proceed until only 100 LNPs remain to compare the polydispersity at the same average particle volume.

### Charge regulation and surrogate model

The potentials *ψ* in the DLVO equation (Eq. 3) depend on the exact composition and size of the LNP and must be calculated by solving the charge regulation relations^51^. Using the Donnan approximation^52^ we approximate the potential *ψ* anywhere inside the LNP is a constant and equal to the surface potential *ψ*_0_. Thus, the charge density is,

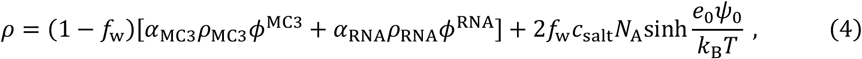

Here *f*_w_ is the water volume fraction, ρ_MC3_ and ρ_RNA_ are the charge density of cationic lipid and siRNA, α_MC3_and α_RNA_ are the degree of ionizations, *ϕ*^MC3^and *ϕ*^RNA^ are volume fractions, *N*_A_ is the Avogadro number, and *e*_0_ is the elementary charge. The surface potential depends on the charge density ρ as:

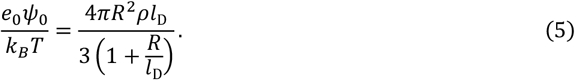

A root-finding approach is used to determine the charge density that satisfies the charge regulation equations (Eqs. 4 and 5). To make the model usable for all salt concentration, a surrogate neural network regression model is trained with data varying nanoparticle sizes, RNA fractions, and salt concentrations. The model efficiently predicts surface potential across a broad parameter space (see Figures S5 and S6). This trained data is provided as a part of the FormLNP kMC code and predictive toolbox.

### Feature importance calculations

A random forest regression model was implemented to analyze the influence of formulation parameters on LNP size and the ratio of empty LNPs. The input variables included initial particle size, PEG MW, PEG fraction, and salt concentration, while the output variables were LNP size and the ratio of empty LNPs obtained from kMC simulations. Prior to training, all input variables were standardized. Separate random forest models were trained for each output variable (LNP final size, ratio of empty LNPs and payload distribution variance). The models were evaluated using 5-fold cross-validation with root mean squared error (RMSE) as the performance metric. Feature importance was determined using the mean decrease in impurity (Gini importance) from the trained models. All analyses, including data preprocessing, model training, cross-validation, and visualization, were conducted in Python using the scikit-learn machine learning package.

## Associated content

### Data Availability Statement

The data supporting the findings of this study are available within the Article and its Supplementary Information. All data are available from the corresponding author upon request.

## Code availability

The “FormLNP” computational framework that predicts LNP size and RNA payload distribution can be found on this website: https://sites.google.com/view/formlnp/home

All other custom scripts and tools used in this study are available from the corresponding author upon request.

## Supporting Information

The Supporting Information includes detailed experimental procedures, molecular dynamics (MD) simulation parameters, kinetic Monte Carlo (kMC) simulation methodology^34,53^, and machine learning approaches for charge regulation modeling. It also contains details of the predictive toolbox for LNP size and siRNA payload distribution.

## Acknowledgement

This work is supported by start-up funds provided by the Whiting School of Engineering at JHU to TC, a grant from the National Cancer Institute to HQM (R01CA293906-01A1), and the National Institute of Allergy and Infectious Diseases (R01AI183336, R01AI181217) to THW. Computational work was carried out at the Advanced Research Computing at Hopkins (ARCH) core facility (rockfish.jhu.edu), which is supported by the National Science Foundation (NSF) grant number OAC 1920103.

