## Supporting Information for "Controlling Payload Heterogeneity in Lipid Nanoparticles for RNA-Based Therapeutics"

Turash Haque Pial[1*], Sixuan Li[2], Jinghan Lin[1,3,5], Tza-Huei Wang[2,3,4], Hai-Quan Mao[1,3,4,5], Tine Curk[1,3*]
[1]Department of Materials Science and Engineering, Johns Hopkins University, Baltimore
[2]Department of Mechanical Engineering, Johns Hopkins University, Baltimore
[3]Institute for NanoBioTechnology, Johns Hopkins University, Baltimore
[4]Department of Biomedical Engineering, Johns Hopkins University School of Medicine, Baltimore
[5]Translational Tissue Engineering Center, Johns Hopkins University School of Medicine, Baltimore

This supplementary information contains details of the experimental setup (Figure S1), molecular dynamics simulation details and supporting data (Figures S2-S4), Monte Carlo simulations and surrogate model for electrostatics (Figures S5-S7), theoretical analysis of merging barriers (Figure S8) and SHAP analysis (Figure S9).

### S1. Single particle analysis of siRNA LNP on CICS

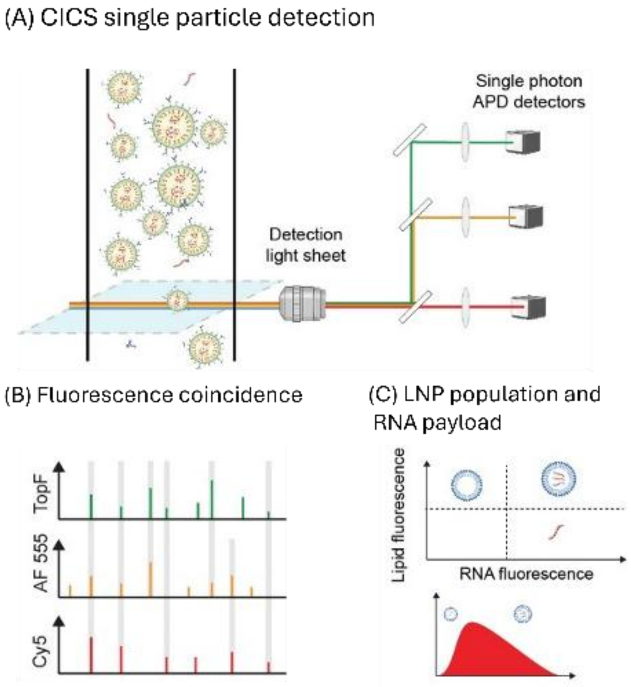


***Figure S1***. *CICS single nanoparticle detection and analysis of siRNA-LNP. (A) The fluorescently labeled LNP sample was applied into the microfluidic flow channel and detected by the cylindrical illumination (light sheet) three-color lasers. The florescence of each event (including LNPs and unencapsulated siRNAs) was collected by the single photon counting APD detectors after the confocal configuration. (B) The three-color fluorescence and their coincidence were analyzed for the sample population classification. (C) Each fluorescence event was classified into siRNA encapsulated LNPs, empty LNPs and unencapsulated siRNAs. For the siRNA encapsulated LNP population, the fluorescence distribution was analyzed by a deconvolution algorithm to obtain the siRNA payload distribution at single nanoparticle level.*

### S2. Molecular dynamics simulation

We employed the Cooke and Deserno 3-bead lipid model[1]. In this model, a lipid consists of one "head" bead and two "tail" beads, with bead sizes fixed by the Weeks-Chandler-Andersen (WCA) potential:

$$V(r,b) = \begin{cases} 4\epsilon_{LJ}[(b/r)^{12} - (b/r)^6 + 1/4], & r \le r_c \\ 0, & r > r_c \end{cases}$$

We used $\sigma = 0.72\ \mathrm{nm}$ as the unit of length and $\epsilon_{LJ} = 1$ set the energy scale. $b_{head,head} = b_{head,tail} = 0.95\sigma$ and $b_{tail,tail} = \sigma$ ensure a cylindrical lipid shape. The cutoff for all repulsive interactions is fixed at $r_c = 2^{1/6}b$, ensuring purely repulsive interactions among beads except modification mentioned below. Three beads of the lipids are connected by two finite extensible nonlinear elastic (FENE) bonds. Lipids are straightened by a harmonic spring with rest length $4\sigma$ between head bead and second tail bead. We also add attractive potential in tail-tail interactions addressing hydrophobic effect (see the equation below). Parameters of these angles, and tail-tail interactions are similar to the original model.[1]

The original Cooke and Deserno model was developed to simulate lipid bilayers and does not facilitate the spontaneous formation of nanoparticles, even in the presence of siRNA. This limitation arises because the model includes attractive interactions only between tail beads, while interactions between head beads are purely repulsive. As a result, lipids preferentially assemble into planar bilayers, with no driving force for forming nanoparticles. We introduced a short-range, weak attractive interaction between head beads. This interaction captures effective solvent-mediated attractions. We used the same cosine-squared potential form as employed for tail–tail attraction in the original model, ensuring consistency. The new head–head potential is defined as:

$$V(r) = \begin{cases} -\epsilon_a, & r < r_c \\ -\epsilon_a cos^2 \dfrac{\pi(r - r_c)}{2w_c}, & r_c \le r \le r_c + w_c \\ 0, & r > r_c + w_c \end{cases}$$

Potential well depth or strength $\epsilon_a$ and decay range $w_c$ are the key tuning parameters in our model. $r_c$ is kept consistent with other interactions in the system.

To identify parameter sets conducive to nanoparticle formation, we systematically varied $\epsilon_a$ and $r_c + w_c$ (Figure S2). We found that very small values of interaction strength and cutoff distance were insufficient to drive spherical nanoparticle assembly, instead forming bilayers and disk-like structures. Conversely, very large values resulted in a stacked disk-like structure, likely due to the loss of fluidity from strong $\epsilon_a$ and long-range ordering induced by a large $r_c + w_c$. Our simulation shows that, at intermediate ranges, the interaction between lipid heads is strong and fluid enough for them to come close together with RNAs, leading to the formation of spherical nanoparticles. Based on these findings, we used $\epsilon_a$=0.5 and $r_c + w_c = 2.5$ for the primary simulations presented in the manuscript.

siRNA was modeled as a bead–spring polymer, where each coarse-grained bead represents a single RNA base pair. Beads interact via excluded volume repulsion using the WCA potential. Adjacent beads are connected via a harmonic bond potential with an equilibrium bond length of 0.42σ (≈0.3 nm, consistent with base pair rise assuming σ = 0.72 nm). To capture the intrinsic rigidity of RNA, an angular potential was applied to enforce a high persistence length (~50 nm), in agreement with the known stiffness of double-stranded RNA in solution. Each RNA bead was assigned to a mass 4.5 times that of a lipid bead to reflect its higher molecular weight. Polyethylene glycol (PEG) chains were also modeled as bead–spring polymers, with each bead representing a monomer unit. Bonds were defined using the same harmonic potential as RNA. However, no angular potential was included for PEG, reflecting its flexibility at the coarse-grained scale, as its persistence length is approximately equal to its Kuhn length (comparable to the bead size). The mass of each PEG bead was set equal to that of a lipid bead. One end of each PEG chain was covalently linked to an uncharged lipid head group, forming a PEGylated lipid molecule.

Electrostatic interaction between charged beads (negatively charged RNA and positively charged lipid head) was modeled using the Debye–Hückel screened Coulomb potential,

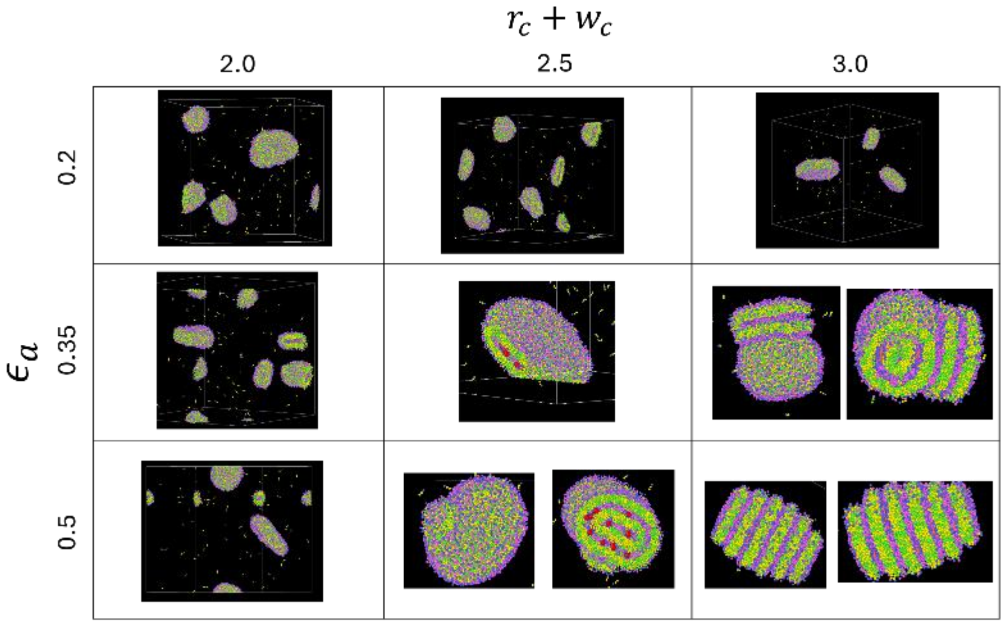


***Figure S2***. *Coarse-grained molecular dynamics (CG-MD) simulations illustrate the impact of cutoff distance ($r_c + w_c$ ) and interaction strength ($\epsilon_a$) on the self-assembly of lipids and RNA. Red beads represent RNA, purple beads correspond to lipid head groups, and green beads denote lipid tail groups.*

which accounts for ionic screening due to salt ions in solution. The potential energy between two charged beads i and j separated by distance r is given by:

$$V_{DH}(r) = \frac{q_i q_j}{4\pi\epsilon r} e^{-k_{DH} r}$$

where: $q_i$, $q_j$ are the charges on the beads, $\epsilon$ is the permittivity of solvent and ${k_{DH}}^{-1}$ is the Debye screening length. In our simulations, the solvent mixture reflects a 3:1 aqueous-to-lipid ratio (we used reduced value of $\epsilon$, corresponding to approximately 25% ethanol in water) and the screening length was calculated assuming a 25 mM concentration of monovalent salt. Other lipid and PEG beads were uncharged and thus did not contribute to electrostatic interactions.

A Langevin thermostat was employed, with a scaling factor introduced in the damping coefficient to ensure proper diffusion of lipids and siRNA in the implicit solvent. This scaling factor was determined by calculating the relative diffusion coefficients of siRNA and lipids using the Stokes-Einstein relation:

$$\frac{D_{lipid}}{D_{RNA}} = \frac{k_B T}{6\pi\eta R_{lipid}} \Big/ \frac{k_B T}{6\pi\eta R_{RNA}} = \frac{R_{RNA}}{R_{lipid}} = \frac{{M_{RNA}}^{1/3}}{{M_{lipid}}^{1/3}} = \frac{13000^{1/3}}{580^{1/3}} \cong 2.8$$

*D* is the diffusion coefficient, *R* is the effective hydrodynamic radius, *M* is the mass, and $\eta$ is the viscosity. The siRNA mass was approximated as 13,000 Da, while the lipid mass was averaged at 580 Da, based on a typical formulation consisting of ionizable lipids (DLin-MC3-DMA , MW 690 Da, 50%), cholesterol (MW 390 Da, 40%), and DSPC (MW 790 Da, 10%).

Although the Stokes–Einstein relation assumes spherical particles, siRNA is better modeled as a short, rigid cylinder with length $L_D$=7 nm (3 nm for lipid) and diameter $d_D$=2 nm (same

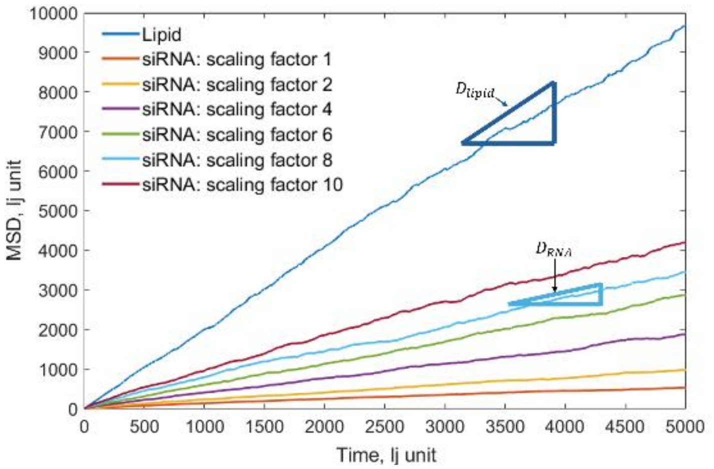


***Figure S3**. Mean square displacement of lipid and siRNA, siRNA damping scaling factor is varied from 1 to 10.*

for lipid). Using the analytical expressions derived by Tirado, Martínez, and de la Torre[2] for cylindrical diffusion in viscous fluids, the orientation-averaged translational diffusion coefficient is:

$$D = \frac{kT}{3\pi\eta L_D} \cdot \frac{1}{\ln\left(\frac{L_D}{d_D}\right) + c1}$$

where $c1$ is an empirical correction for short rods provided in ref. 2. Using above mentioned values of $L_D$ and $d_D$, we can calculate that lipid diffuses approximately 2.4 times faster than the siRNA. This diffusion ratio closely matches the mass-based scaling factor of ~2.8, supporting the validity of using $M^{1/3}$, especially in implicit solvent simulations where full hydrodynamic interactions are absent and the primary goal is to preserve relative mobility between species.

To determine the appropriate scaling factor, we separately simulated the diffusion of lipids and siRNA at very dilute concentrations using a Langevin thermostat. We analyzed the mean squared displacement (MSD) over time, where the slope of MSD in the linear regime provides the diffusion coefficient. By systematically varying the Langevin damping scaling factor (which modifies the frictional drag and thereby affects the diffusion rate), we identified that a scaling factor of 8 yielded a close match to the expected diffusion ratio between the two species; $\frac{D_{lipid}}{D_{RNA}} \cong 2.75$ (Figure S3).

To check the proximity of RNA and lipid near a lipid (as shown in Figure 2B in the main manuscript), we have checked radial distribution function (RDF) of lipid–lipid pairs. A clear first peak followed by a distinct valley is observed around 1.7 LJ units (Figure S4), which we selected as the cutoff distance for coordination number calculations.

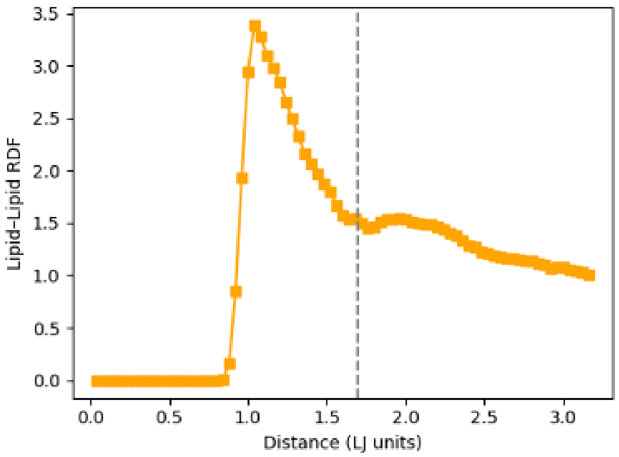


***Figure S4***. *Lipid-lipid radial distribution function.*

### S3. Kinetic Monte Carlo simulations

<u>*Surrogate model for charge regulation:*</u>

The calculation method for surface potential is described in the main manuscript. A machine learning model was trained on synthetically generated data by systematically sampling a 3D parameter space of nanoparticle size, RNA volume fraction ($\phi^{\mathrm{RNA}}$), and salt concentration. Specifically, the training dataset consisted of: radii sampled uniformly from 1 to 20 nm (20 points), RNA volume fractions from 0 to 1 (20 points), salt concentrations from 5 mM to 150 mM (20 points), yielding a total of 8,000 training samples (20 × 20 × 20). Each sample was labeled by numerically solving the nonlinear charge regulation equation under the corresponding conditions to obtain the surface potential. Figure S5 presents a comparison between the surface potential obtained by direct numerical solution and the predictions from the trained neural network model. The agreement is excellent across the entire parameter space. Minor discrepancies occur only at radii below 2 nm, a regime that is not relevant to our system, as all lipid nanoparticles in this study exceed that size.

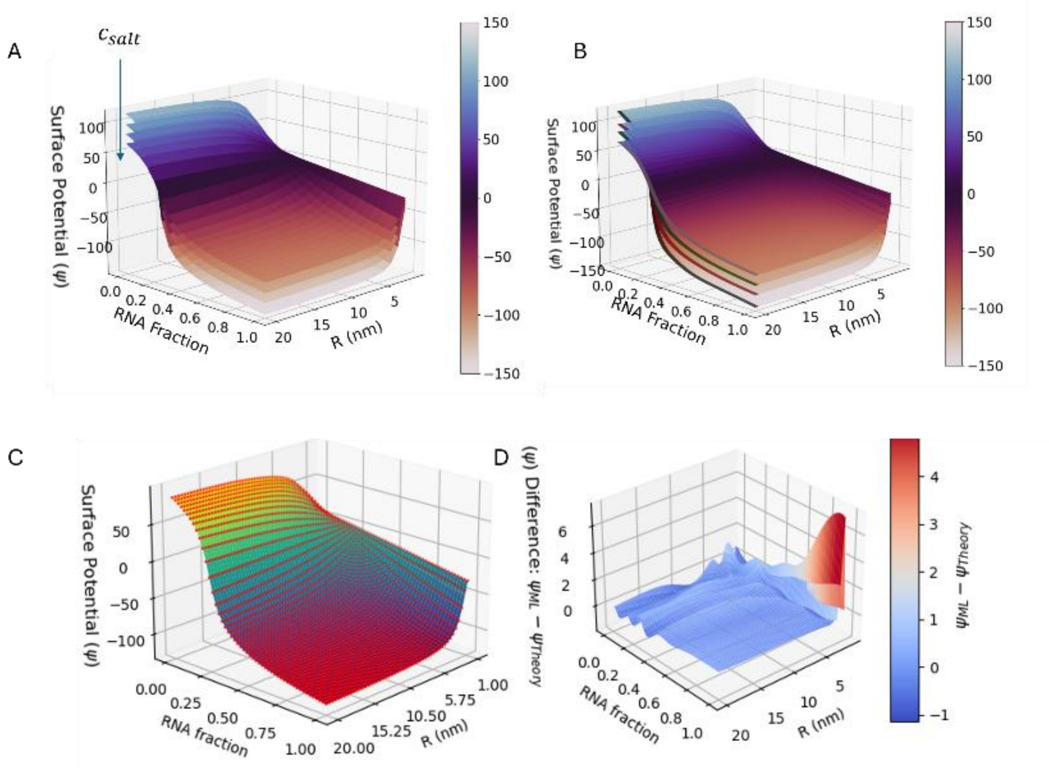


***Figure S5.*** *(A) Numerically solved surface potential (in mV unit), and (B) machine learning obtains surface potential. Color bars indicate $\psi$. The four different surfaces correspond to four different salt concentrations (0.01, 0.025, 0.05 and 0.1 M). Increasing the salt concentration reduces the magnitude of $\psi$. (C) Overlay of predicted surface (smooth surface) and numerically solved values (red dots) for 0.025 M salt. The surface is shown to guide the eye. (D) Difference between two methods for 0.025 M salt case.*

In the machine larning model, we employed a feedforward neural network implemented using the MLPRegressor from scikit-learn library in Python to predict the surface potential. The architecture consisted of three hidden layers, each containing 20 neurons, and used the logistic activation function. The model was trained using a constant learning rate of 0.005. The model was trained on 80% of the data, with 20% held out as an independent test set with a fixed random seed for reproducibility. An internal validation fraction of 0.2 was used to monitor overfitting during training. Early stopping was enabled, halting training when validation loss did not improve for 10 consecutive iterations. The model converged within 1000 iterations in all cases. Parameters of the machine learning architecture were optimized via a grid search over multiple hyperparameter combinations. Specifically, we varied the number and size of hidden layers and tested learning rates of 0.005 and 0.01.

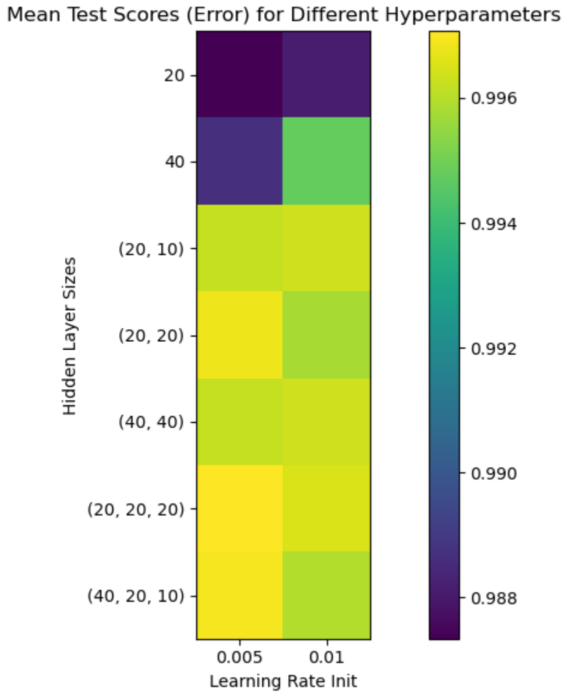


***Figure S6.*** *Hyperparameter optimization.*

<u>*Mean-field theory for LNP growth without RNA:*</u>

The size of LNPs at time $\tau_{\mathrm{M}}$ can be estimated by assuming that for t<$\tau_{\mathrm{M}}$, LNPs can grow in a monodisperse manner without RNA, whereas for time t>$\tau_{\mathrm{M}}$, LNPs grow in the presence of RNA. Initial LNP growth is modeled as a merging rate equation, where integration over time provides the particle size at $\tau_{\mathrm{M}}$. This size serves as the input ($R_0$) for kMC simulations, corresponding to a given mixing flow rate.

The merging rate per particle depends on concentration $c(R)$, diffusivity $D(R)$, and the Arrhenius factor. We can express the time required for a particle to diffuse its radius as $t_D = R^2/(6D(R))$. Using $V_1$ for the average volume of a lipid nanoparticle, we get the rate as:

$$k_m(R) = \frac{2c(R)V_1}{t_D} e^{-E_\mathrm{b}(R)/kT}$$

where $E_\mathrm{b}(R)$ is the fusion energy barrier, calculated as the summation of PEG-PEG repulsion and DLVO interactions, in the absence of RNA. The concentration is given by $c(R) = \frac{c_0 v_o}{V_1}$, where $c_0$ is the lipid concentration and $v_o$ is the volume per lipid, and prefactor 2 accounts for both particles diffusing. Using Stokes-Einstein relation, $D(R) = k_B T/(6\pi\eta R)$, we obtain:

$$k_m(R) = \frac{2k_B T c_0 v_0}{\pi\eta R^3} e^{-E(R)/kT}$$

The evolution of the average particle size, disregarding the polydispersity, is obtained by considering that each merging event between two particles of volume $V_1$ increases the volume per particle by $V_1$, thus,

$$\frac{\partial V_1}{dt} = k_m(R)\frac{V_1}{2}$$

The factor of 2 arises due to stoichiometry since two small particles merge into a single, large one. Assuming LNPs are fully loaded and volume-conserving, the evolution of the average size follows:

$$\frac{\partial R}{dt} = \frac{k_B T c_0 v_0}{3\pi\eta R^2} e^{-E(R)/kT}$$

$$\frac{\partial R}{dt} = k_1 \frac{e^{-(k_2 R^2 + W(R))}}{R^2}$$

Here, $k_1$ tells the merging rate caused by diffusion with no energy barrier:

$$k_1 = \frac{k_B T c_0 v_0}{3\pi\eta},$$

$k_2$ represent the merging barrier for PEG:

$$k_2 = \frac{2\pi c_{PEG} R_F}{3 c_0 v_0}$$

This can be obtained from equation 2 in the main manuscript by considering $\phi_i^{\mathrm{PEG}} = \phi_j^{\mathrm{PEG}} = c_{PEG}/c_0$ and neglecting RNA and water volumes. And $W(R)$ is the DLVO interaction, calculated Eq. 3 in the main manuscript. Here we assumed siRNA fraction ($\phi^{\mathrm{RNA}}$) is zero.

While theoretical models assume spherical nanoparticles, MD simulations revealed a distribution of morphologies, including spherical and disk-like structures. To facilitate consistent comparison with the mean-field model, we computed the mass of each LNP in the MD trajectory and converted it to an equivalent spherical radius. This yielded strong agreement between MD and theory, though the slightly slower growth observed in MD. The slight deviation is caused by the lack of hydrodynamic interaction in MD simulations, which would increase diffusion of assembled particles.

To convert the empirical relationship between flow rate and mixing timescale, we used Ref. 26 and [3] of the main manuscript, where the equation was calibrated using linear PEI (lPEI) with a molecular weight of 22 kDa and a diffusivity of $1\times 10^{-6}\ \mathrm{cm}^2/s$. According to data from fluorescence correlation spectroscopy[4], free siRNA in buffer at has a diffusivity of approximately $1.75\times 10^{-6}\ \mathrm{cm}^2/s$, yielding a diffusivity ratio of 1.75.

<u>*Coalescence rate kernel analysis:*</u>

The ratio $K_{v_1,v_2}/\ K_{v/2,v/2}$ tells the rate of coalescence between two LNPs of different sizes (volumes $v_1$ and $v_2$), relative to two LNPs of equal size (volume $\frac{v}{2}$) merging to form a larger particle of volume $v$. For these calculations, we assume our final LNP has a radius of 20 nm. We divided the total volume $v$ into 20 pairs of smaller LNPs, each of which can merge with another to form a larger particle of volume $v$.

A coalescence kernel consists of two main elements: the diffusion of particles and the energy barrier to merging. The energy barrier is influenced by two factors: radius-dependent interactions, which depend on the size of the particles and affect the likelihood of coalescence, and radius-independent factors, which include environmental influences and other constants that can be assumed to size-independent. To account for these radius-independent factors, we introduced a prefactor that incorporates environmental and other factors affecting coalescence. By combining this prefactor with diffusion and radius-dependent interactions, we captured the full coalescence behavior.

Perfector for PEG-induced barrier (cf. Eq. 2 in the main manuscript) is calculated as:

$$C_{PEG} = \frac{2\pi R_{\mathrm{F}}}{3v_0(1-f_w)}\phi_i^{PEG}$$

Using typical values $(R_{\mathrm{F}} = 3.6, v_0 = 3\ nm^3, f_w = 0.2, and\ \phi_i^{PEG} = 0.015)$, we get $C_{PEG} = 0.047$.

For pH=4 and salt concentration of 0.025 M; surface potential reaches around 80 mV [Figure S7(A)]. As it is plateauing after 10 nm, we assume a constant surface potential at 80 mV. This simplifies the DLVO equation to:

$$W(d,\ R_1, R_2) = -\frac{AR_1R_2}{6d(R_1+R_2)} + \epsilon\frac{R_1R_2\psi^2}{2(R_1+R_2)}\log\left(1+e^{-\frac{d}{l_D}}\right)$$

We can try to isolate the prefactor by

$$C_{\mathrm{DLVO}} = W(d,\ R_1, R_2)/(\frac{R_1R_2}{(R_1+R_2)})$$

For pH=4 and salt concentration of 0.025 M, using constant surface potential, we have found a $C_{\mathrm{DLVO}}$ =0.415.

These values of $C_{PEG}$ and $C_{DLVO}$ is used in Figure 5B in the main manuscript.

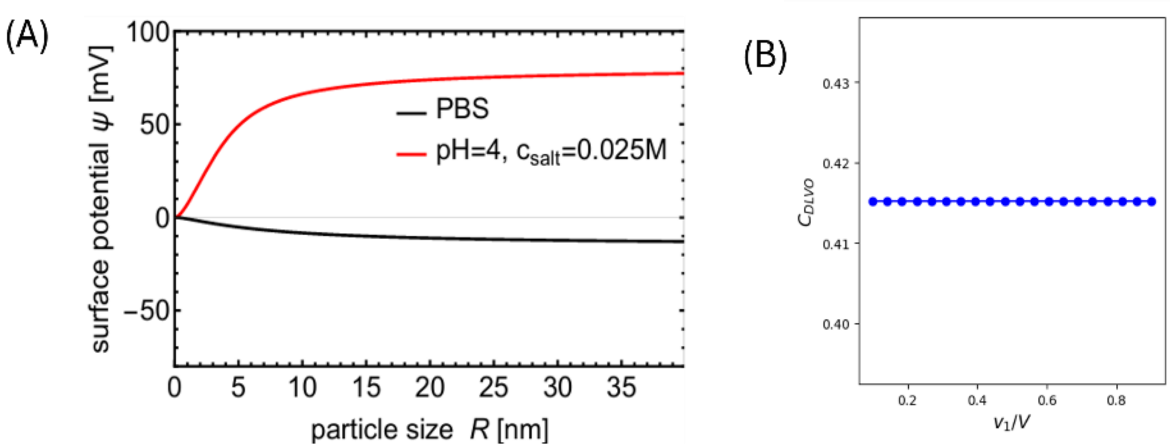


***Figure S7***: *(A) At pH 4 and a salt concentration of 0.025 M, the surface potential is approximately 80 mV for LNPs larger than 10 nm. In PBS, the surface potential is very small. (B)* $C_{DLVO}$ *value obtained for the assumed surface potential of 80 mV.*

In Figure 8, we discuss the relative effect of $C_{PEG}$ and $C_{DLVO}$ in LNP merging process. For PBS cases (after dialysis), energy barrier should be lower as the salt concentration increases, and surface potential is reduced. As a result, we will get smaller $C_{\mathrm{DLVO}}$, leading to a reduced influence of DLVO on the merging rate, and the process will increasingly resemble diffusion-limited merging. In Figure S8 (left), we plot the variation of $C_{\mathrm{DLVO}}$ which indicates the effect of salt concentration on probability of coalescence.

It is also instructive to compare how the functional dependence on particle radius influences merging rates under different interaction barriers. In particular, the DLVO energy barrier scales as $\frac{R_1 R_2}{R_1+R_2}$, while the PEG–PEG steric repulsion barrier scales as $R_1 R_2$. To isolate the

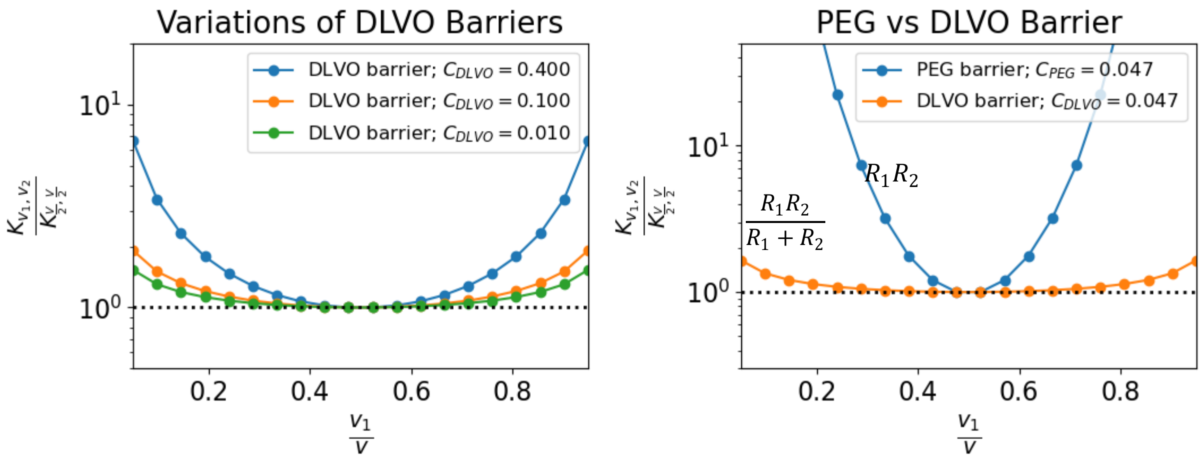


***Figure S8****: (Left) Effect of $C_{DLVO}$ in the probability of coalescence. Smaller $C_{DLVO}$ will result in lesser effect on the probability of coalescence. (Right) PEG barrier will result in higher probability of merging unequal LNPs, even with similar prefactor values to DLVO; showcasing the squared exponential behavior of PEG compared to exponential behavior of DLVO.*

effect of this radius dependence, we performed calculations using the same prefactor for both interactions ($C_{PEG} = C_{DLVO} = 0.047$) but applied their respective barrier functions, as shown in Figure S7 (right). Despite the identical prefactor, the results reveal that merging remains highly selective when only PEG–PEG interactions are considered, whereas the merging rate is significantly suppressed under the DLVO-only model. This contrast highlights the critical role of radius scaling in determining the energy barrier and thus the kinetic accessibility of merging events.

*SHapley Additive exPlanations (SHAP):*

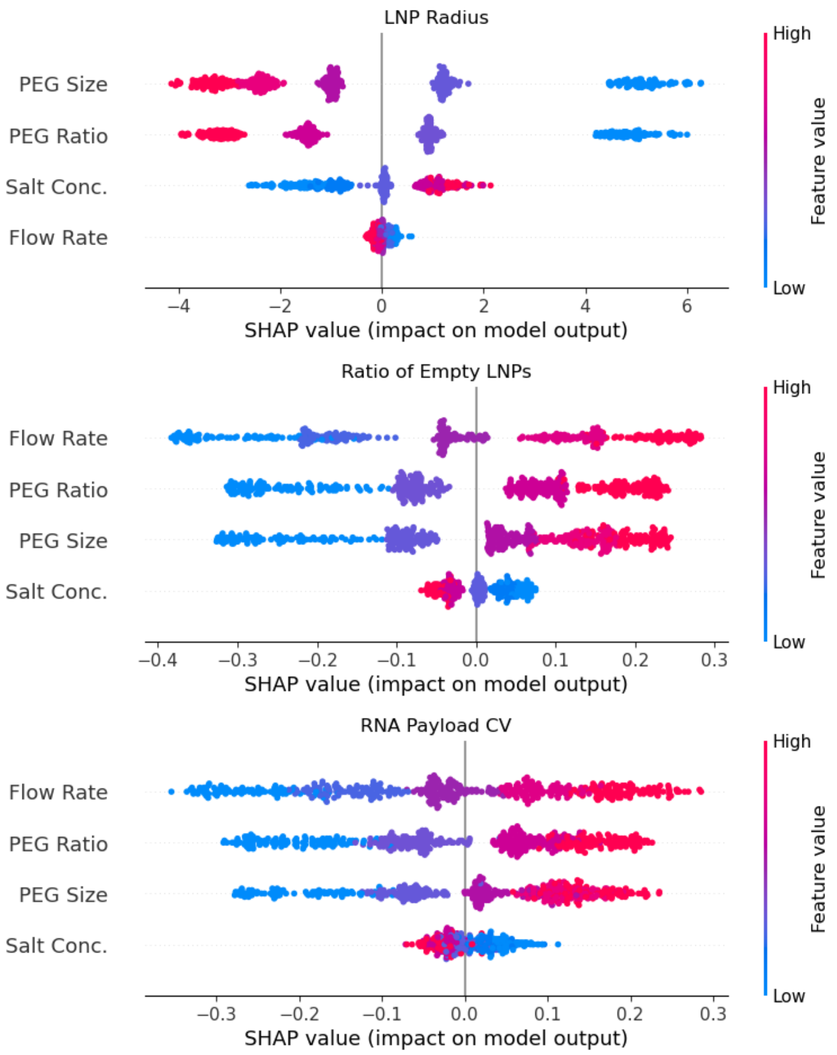


**Figure S9:** *Corresponding SHapley Additive exPlanations (SHAP) values of the feature importance shown in Figure 6A.*

### S4. Computational Toolbox

We have shared an open-source computational toolbox alongside this manuscript to support predictive modeling and kinetic analysis of RNA–LNP systems. This toolbox can be accessed from this website and can be easily used using google-colab which doesn't require any installation.  The toolbox consists of two key components:

Predictive Modeling Module: This includes trained machine learning models and prediction scripts that allow users to estimate key LNP characteristics, including final size, empty LNP ratio, and RNA payload distribution variation; based on formulation parameters such as mixing flow rate, salt concentration, PEG molecular weight, and PEG/lipid ratio. This module is designed for users seeking rapid prediction and screening of formulation outcomes without requiring additional simulation.

kMC Simulation Module: For users interested in mechanistic insight and customizable modeling, we provide the full kMC simulation framework. This component includes the charge regulation-based surface potential model, and simulation scripts for LNP growth dynamics. Users can modify these scripts to explore specific kinetic regimes, observe LNP size evolution, and monitor empty particle fractions over time.

Interested readers can access the full toolbox, including all scripts, trained models, and usage instructions, at this website:

https://sites.google.com/view/formlnp
